# The Integrated Rapid Infectious Disease Analysis (IRIDA) Platform

**DOI:** 10.1101/381830

**Authors:** Thomas C Matthews, Franklin R Bristow, Emma J Griffiths, Aaron Petkau, Josh Adam, Damion Dooley, Peter Kruczkiewicz, John Curatcha, Jennifer Cabral, Dan Fornika, Geoffrey L. Winsor, Melanie Courtot, Claire Bertelli, Ataollah Roudgar, Pedro Feijao, Philip Mabon, Eric Enns, Joel Thiessen, Alexander Keddy, Judith Isaac-Renton, Jennifer L. Gardy, Patrick Tang, The IRIDA Consortium João A Carriço, Leonid Chindelevitch, Cedric Chauve, Morag R Graham, Andrew G McArthur, Eduardo N Taboada, Robert G Beiko, Fiona SL Brinkman, William WL Hsiao, Gary Van Domselaar

## Abstract

Whole genome sequencing (WGS) is a powerful tool for public health infectious disease investigations owing to its higher resolution, greater efficiency, and cost-effectiveness over traditional genotyping methods. Implementation of WGS in routine public health microbiology laboratories is impeded by a lack of user-friendly automated and semi-automated pipelines, restrictive jurisdictional data sharing policies, and the proliferation of non-interoperable analytical and reporting systems. To address these issues, we developed the Integrated Rapid Infectious Disease Analysis (IRIDA) platform (irida.ca), a user-friendly, decentralized, open-source bioinformatics and analytical web platform to support real-time infectious disease outbreak investigations using WGS data. Instances can be independently installed on local high-performance computing infrastructure, enabling private and secure data management and analyses according to organizational policies and governance. IRIDA’s data management capabilities enable secure upload, storage and sharing of all WGS data and metadata. The core platform currently includes pipelines for quality control, assembly, annotation, variant detection, phylogenetic analysis, *in silico* serotyping, multi-locus sequence typing, and genome distance calculation. Analysis pipeline results can be visualized within the platform through dynamic line lists and integrated phylogenomic clustering for research and discovery, and for enhancing decision-making support and hypothesis generation in epidemiological investigations. Communication and data exchange between instances are provided through customizable access controls. IRIDA complements centralized systems, empowering local analytics and visualizations for genomics-based microbial pathogen investigations. IRIDA is currently transforming the Canadian public health ecosystem and is freely available at https://github.com/phac-nml/irida and www.irida.ca.

**Impact Statement:** Whole genome sequencing (WGS) is revolutionizing infectious disease analysis and surveillance due to its cost effectiveness, utility, and improved analytical power. To date, no “one-size-fits-all” genomics platform has been universally adopted, owing to differences in national (and regional) health information systems, data sharing policies, computational infrastructures, lack of interoperability and prohibitive costs. The Integrated Rapid Infectious Disease Analysis (IRIDA) platform is a user-friendly, decentralized, open-source bioinformatics and analytical web platform developed to support real-time infectious disease outbreak investigations using WGS data. IRIDA empowers public health, regulatory and clinical microbiology laboratory personnel to better incorporate WGS technology into routine operations by shielding them from the computational and analytical complexities of big data genomics. IRIDA is now routinely used as part of a validated suite of tools to support outbreak investigations in Canada. While IRIDA was designed to serve the needs of the Canadian public health system, it is generally applicable to any public health and multi-jurisdictional environment. IRIDA enables localized analyses but provides mechanisms and standard outputs to enable data sharing. This approach can help overcome pervasive challenges in real-time global infectious disease surveillance, investigation and control, resulting in faster responses, and ultimately, better public health outcomes.

**DATA SUMMARY:**

1. Data used to generate some of the figures in this manuscript can be found in the NCBI BioProject PRJNA305824.

## INTRODUCTION

Infectious diseases continue to exact a substantial toll on health and health-care resources accounting for nearly a quarter of the estimated 52.8 million deaths annually, as well as hundreds of billions of dollars in lost productivity representing significant percentage of global GDP (1,2). Globalization of food networks increases opportunities for the spread of foodborne pathogens beyond borders and jurisdictions. Furthermore, new foodborne pathogens emerge driven by factors such as pathogen evolution or changes in agricultural and food manufacturing practices (3). In response to these challenges, public health microbiology surveillance programs employ molecular methods for routine typing and monitoring of foodborne pathogens (4–7). Current techniques such as Pulsed-Field Gel Electrophoresis (PFGE) and traditional Multi-Locus Sequence Typing (MLST) offer far less discriminatory power for distinguishing outbreak cases from sporadic cases compared with more recently developed Whole Genome Sequence (WGS)-based approaches such as core genome MLST (cgMLST) or Single Nucleotide Variant/Polymorphism (SNV/SNP)-based phylogenies (8). Furthermore, conventional typing methods are resource intensive and can require relatively long turnaround times. In contrast, WGS can inform investigators about numerous infectious pathogen traits in a single “assay”, often with reduced turnaround time and with improved analytical power. As such, public health and food safety authorities are undergoing a historic transition, as WGS-based applications increasingly replace many molecular and phenotypic assays, ranging from serotyping and other molecular level microbial characterization for pathogen identification and surveillance as well as outbreak response, to antimicrobial resistance (AMR) and virulence prediction for risk assessment (9–17).

While the advantages of implementing WGS-based analytics are widely recognized, efforts to integrate WGS into regional and national foodborne pathogen surveillance programs have been hampered by the lack of easy-to-use and validated bioinformatics tools, particularly impacting implementation in regional public health microbiology laboratories. To date, no “one-size-fits-all” genomics platform has been developed or universally accepted owing to differences in national health systems, data sharing policies, computational infrastructures, prohibitive costs, lack of interoperability, and lack of qualified personnel to carry out analyses. Several international infectious disease monitoring initiatives are working on approaches to tackle these problems. PulseNet (www.pulsenetinternational.org/), a molecular subtyping network for foodborne disease surveillance, is building towards routine implementation of WGS for foodborne surveillance and cluster detection for coordinated outbreak response. PulseNet applies BioNumerics (Applied Maths, Sint-Martens-Latem, Belgium), a widely-used commercial analysis platform that recently has incorporated whole genome sequence assembly and typing functionality into its suite of molecular data analysis, management, and reporting tools. The international GenomeTrakr network is the first distributed network of laboratories applying WGS to coordinate investigations of outbreaks of foodborne illnesses with compliance actions (18). The Global Microbial Identifier (GMI) pathogen tracking initiative collaboratively works with international health, regulatory and research organizations, as well as the International Nucleotide Sequence Database Collaboration (INSDC), to create a worldwide network of shared genomic information for bacterial, viral, and parasitic microorganisms for identification of and response to infectious disease clusters. The National Institutes of Health - National Center for Biotechnology Information (NIH-NCBI) Pathogen Detection pipeline and database offers centralized services for genome assembly, annotation, strain clustering and AMR prediction for foodborne pathogens. Such centralized services for analyses and data management, however, have limitations for public health agencies dealing with sensitive data under jurisdiction of other authorities, which require data and analyses to be performed and maintained locally. Commercial platforms can be extremely costly if multiple licenses are required to outfit multiple laboratories within a broader network. Furthermore, centralized services and proprietary systems have less flexibility than open-source software, resulting in the need for users to create in-house workarounds. As such, a number of in-house tools and platforms, have emerged in parallel, aiming to fill the void and address the distinct needs of different organizations. The Advanced Molecular Detection initiative of the US Centers for Disease Control and Prevention (CDC) supports the building of shared computing and laboratory capacity for genomics, bioinformatics, and next-generation diagnostic testing, and continues to rollout WGS technologies to state public health laboratories. Public Health England has implemented whole genome sequencing as a routine typing tool for public health surveillance of *Salmonella*, and has continued to rollout standardized WGS-based workflows via its Pathogen Genomics Service (15,19). INNUENDO is an integrated genomics-based foodborne pathogens surveillance project developed as a collaborative effort between several European institutions, co-funded by the European Food Safety Authority. The INNUENDO initiative aims to provide a lightweight framework for the integration of bacterial whole genome sequencing in routine surveillance and epidemiological investigations supporting small European countries with limited resources (sites.google.com/site/theinnuendoproject). Similarly, the European COMPARE research network aims to provide an analytical framework and data exchange platform for sequence-based pathogen data, thereby contributing to containment and mitigation of emerging infectious diseases (compare-europe.eu) (20). The WGSA platform (wgsa.net), created by the Centre for Genomic Pathogen surveillance, provides standardized pipelines for users to analyze and visualize genomic assemblies. Nullarbor (Microbiological Diagnostics Unit Public Health Laboratory, University of Melbourne, Australia; github.com/tseemann/nullarbor) and the Center for Genomic Epidemiology (Danish Technical University, Denmark; genomicepidemiology.org), are microbial sequence analysis platforms containing pipelines for the assembly, annotation and analysis of sequenced isolates, and produce public health microbiology reports supporting research and foodborne outbreak investigations.

Similarly, Canada has developed its own frameworks for building bioinformatics and public health genomics capacity according to its policies and strategic goals. Canada’s public health system exists as a decentralized, multi-jurisdictional network of provincial and federal laboratories collectively referred to as the Canadian Public Health Laboratory Network (CPHLN). In 2016, the CPHLN initiated a national public health genomics strategic goal with a special emphasis on accommodating the specific requirements and capabilities of each participating lab while adhering to published best practices. Best practices for implementing WGS in clinical settings for diagnostics of infectious disease include, as essential components for meeting regulatory standards: the provision of documentation for validation and quality assurance, high capacity data storage, version traceability, and data transfer confidentiality (21–23). In addition to clinical best practices, current best practices in bioinformatics for food safety authorities are being created to increase the transparency of food microbiology risk management decision making (24).These practices include the implementation of data standards, maintainability of software, demonstration of data integrity, traceability, and auditability, as well as the development of frameworks for reporting and interpretation. With these best practices in mind, we have developed the Integrated Rapid Infectious Disease Analysis (IRIDA) platform, a secure, decentralized, open-source, freely available, end-to-end public health genomics platform for microbial infectious disease investigations. IRIDA enables organizations to perform localized analyses, while simultaneously supporting data sharing and synchronization across jurisdictions, and sequence deposition to centralized repositories. The IRIDA project was originally developed as a trilateral collaboration between the Public Health Agency of Canada, the British Columbia Centre for Disease Control Public Health Laboratory, and Simon Fraser University, with key contributions from additional academic, federal, and provincial partners, representing national and provincial public health as well as research interests. Thereafter, the project has expanded to include additional collaborations with academic and government partners. Here, we describe the architecture and core functionality of the IRIDA platform, its current usage in Canada, and its emerging uptake by the global community.

## Results

IRIDA is a web-based application that can be locally deployed as a stand-alone platform, designed to provide public health, clinical microbiology, and food regulatory authorities with the capability to incorporate next-generation sequencing into their surveillance, diagnostics, reference typing, and research programs. IRIDA’s core functionality encompasses four main areas: 1) data management, 2) user management and data sharing, 3) data analysis, and 4) reporting and visualization. We describe each of these main functions here.

### Data Management

IRIDA performs all aspects of sequence data and metadata management, including data import, export, storage, organization, tracking, and sharing (Fig. 1).

**Figure 1.**
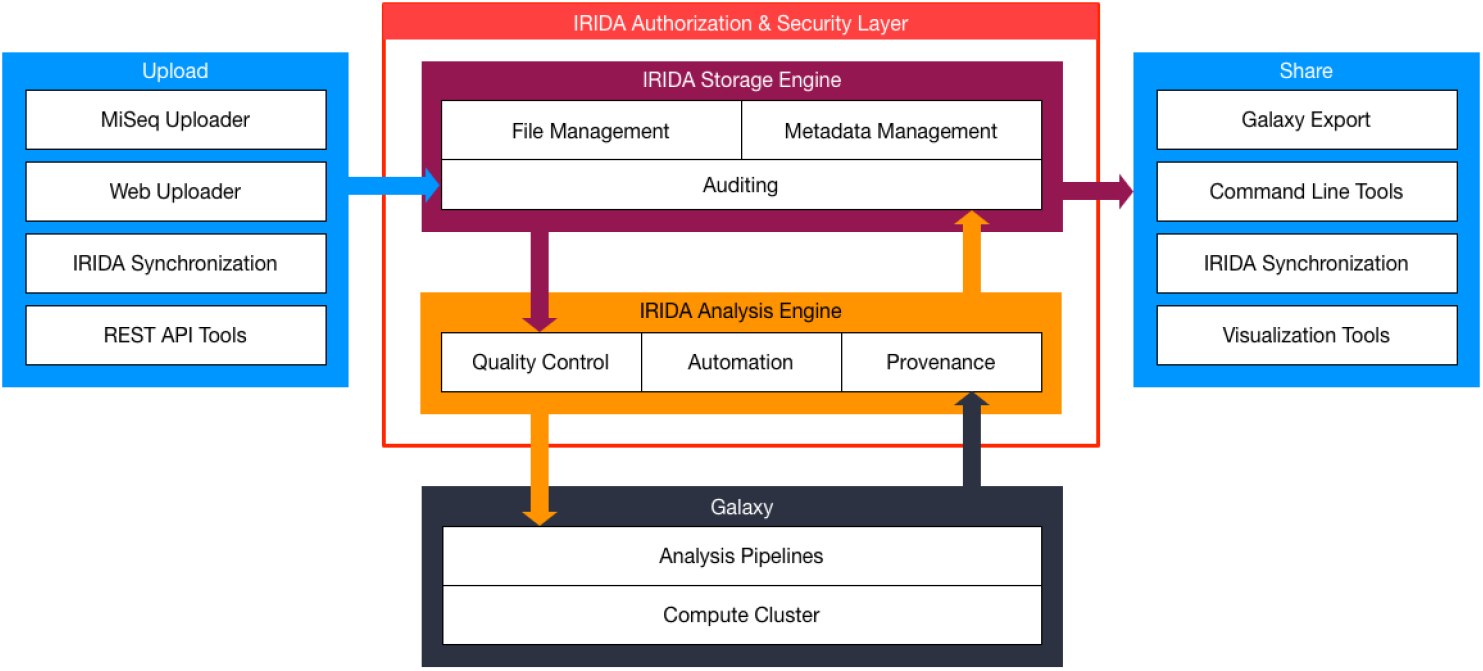
IRIDA platform schematic overview. The IRIDA platform provides an end-to-end solution for sequence data management and analysis. This includes multiple options for importing data into the system including a custom Uploader Tool that runs on an Illumina MiSeq sequencer, a web interface for uploading data, or synchronization from other IRIDA installations. IRIDA’s authorization and security layer validates that a user is able to access or modify the data they are attempting to access. Once authenticated, IRIDA’s auditing layer creates an audit trail for all data uploaded, modified, or deleted. IRIDA contains an analysis engine to perform quality control on all uploaded sequencing data, provides automation tools, and stores data provenance. The analysis engine communicates with Galaxy to schedule and execute its analysis pipeline tools. IRIDA also provides multiple secure data sharing options, ranging from exporting data to Galaxy for further analysis, to command-line tools, to data synchronization between IRIDA installations.

IRIDA’s data structure is rooted by a Project. Projects contain a collection of Samples, which contain a collection of Sequence Data. Projects also contain Analyses generated from the sequence data within that project. Metadata can be associated with Projects, Samples, Sequence Data, and Analyses (Fig. 2). IRIDA’s data structure is modeled after, and fully compatible with, the BioProject data structure used by INSDC databases (NCBI, EMBL-EBI and DDBJ). IRIDA’s data structure facilitates deposition of reads, assemblies, and annotated contigs to the various public sequence repositories.

**Figure 2.**
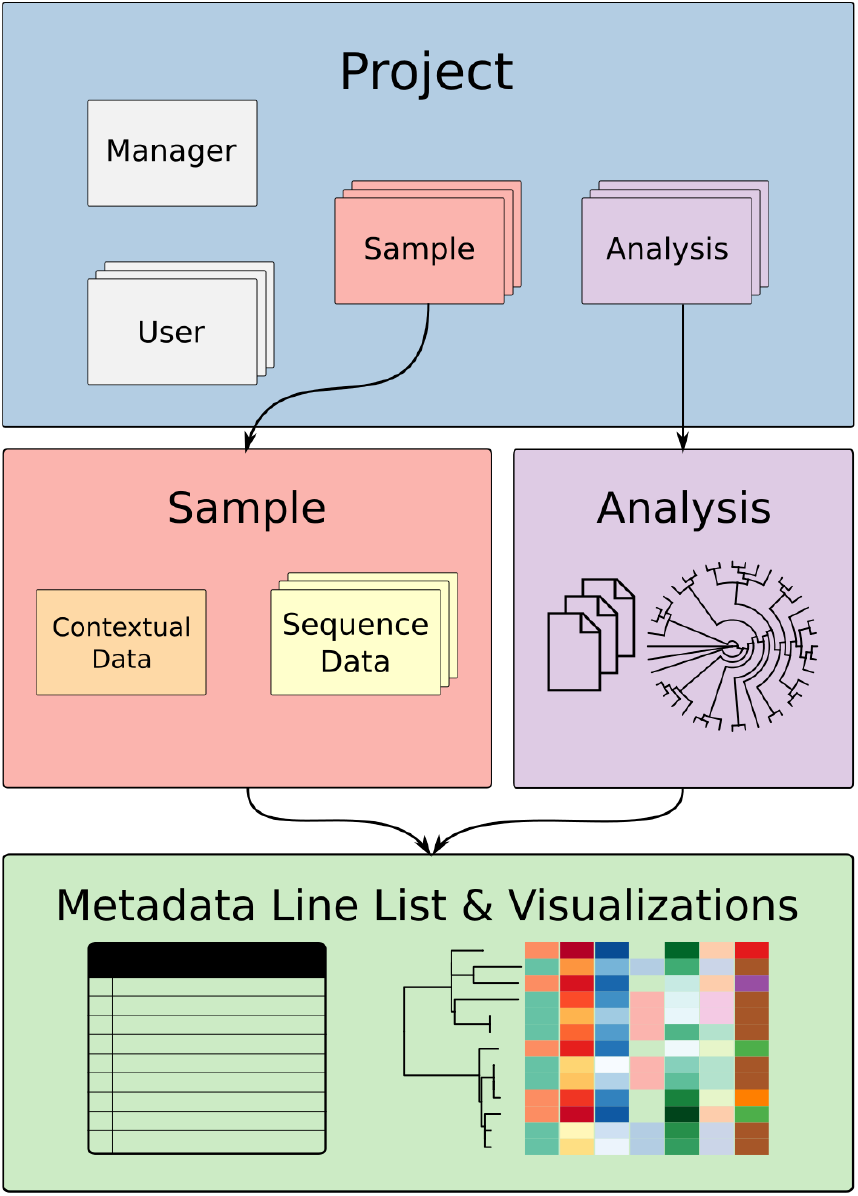
Project structure and data model. IRIDA’s data structure is rooted by a “Project”. A Project is a container for any number of related samples, their sequencing data, and analysis results. Metadata associated with the project, sequences and analyses are also stored within the Project. A “Sample” container stores contextual data (*e.g*. environmental, clinical, food source, collection dates, and geographical locations) alongside sequencing data, better enabling integrated analyses. Data access is provided on a per-project basis. Users who are added to a project are given access to all data contained within the project.

IRIDA currently provides a stand-alone sequence data Uploader Tool for Illumina sequencing instruments; uploaders for other sequencing platforms are under development. The sequence data Uploader Tool monitors the sequencer’s output folders and will automatically import new sequence data and associated metadata into IRIDA. Users can also import sequence data and metadata into IRIDA directly via the web interface. To preserve data integrity, sequence data is stored in IRIDA’s underlying file system with “read-only” access permissions.

IRIDA uses a familiar “online shopping cart” model to assist users in running their analyses: the user selects the data within their projects that they wish to analyze and adds them to the “cart” (Fig. 3). Users then choose their desired analysis pipeline, adjust the pipeline’s runtime parameters (or select a pre-saved set of runtime parameters), and launch their analysis (Fig. 4). IRIDA manages the execution of their analysis and provides indicators for quality control (QC) and execution status.

**Figure 3.**
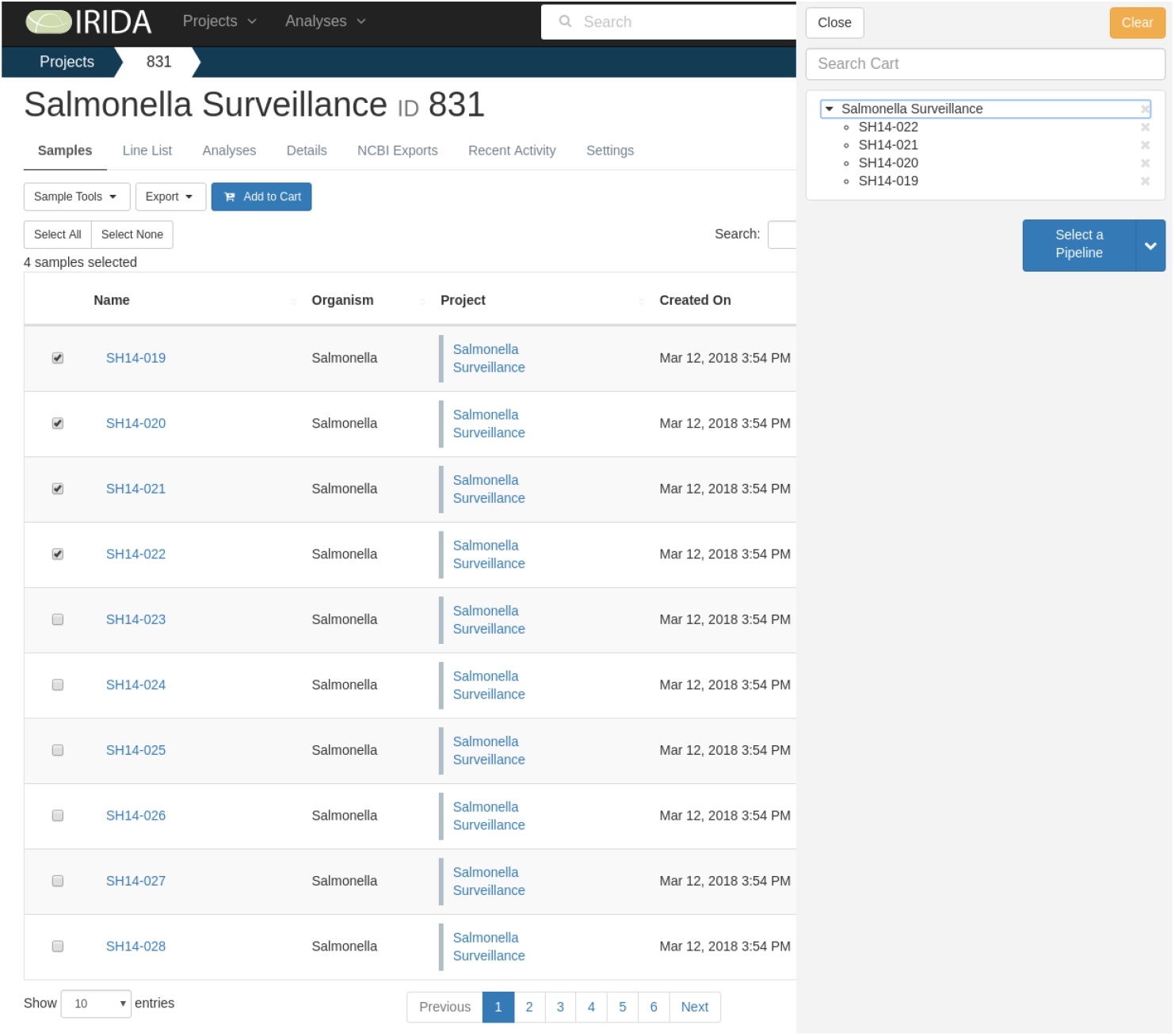
Project Sample page with “Shopping cart” functionality. IRIDA’s project sample page provides point-and-click access to all sample data stored within a project. Sample data can be sorted, filtered, and managed from within this interface. From this page samples can be selected and grouped into IRIDA’s “shopping cart”. The cart provides a familiar means of collecting samples into a temporary storage space for users to later perform further actions. The cart is the entry point for all analysis pipelines in the IRIDA system.

**Figure 4.**
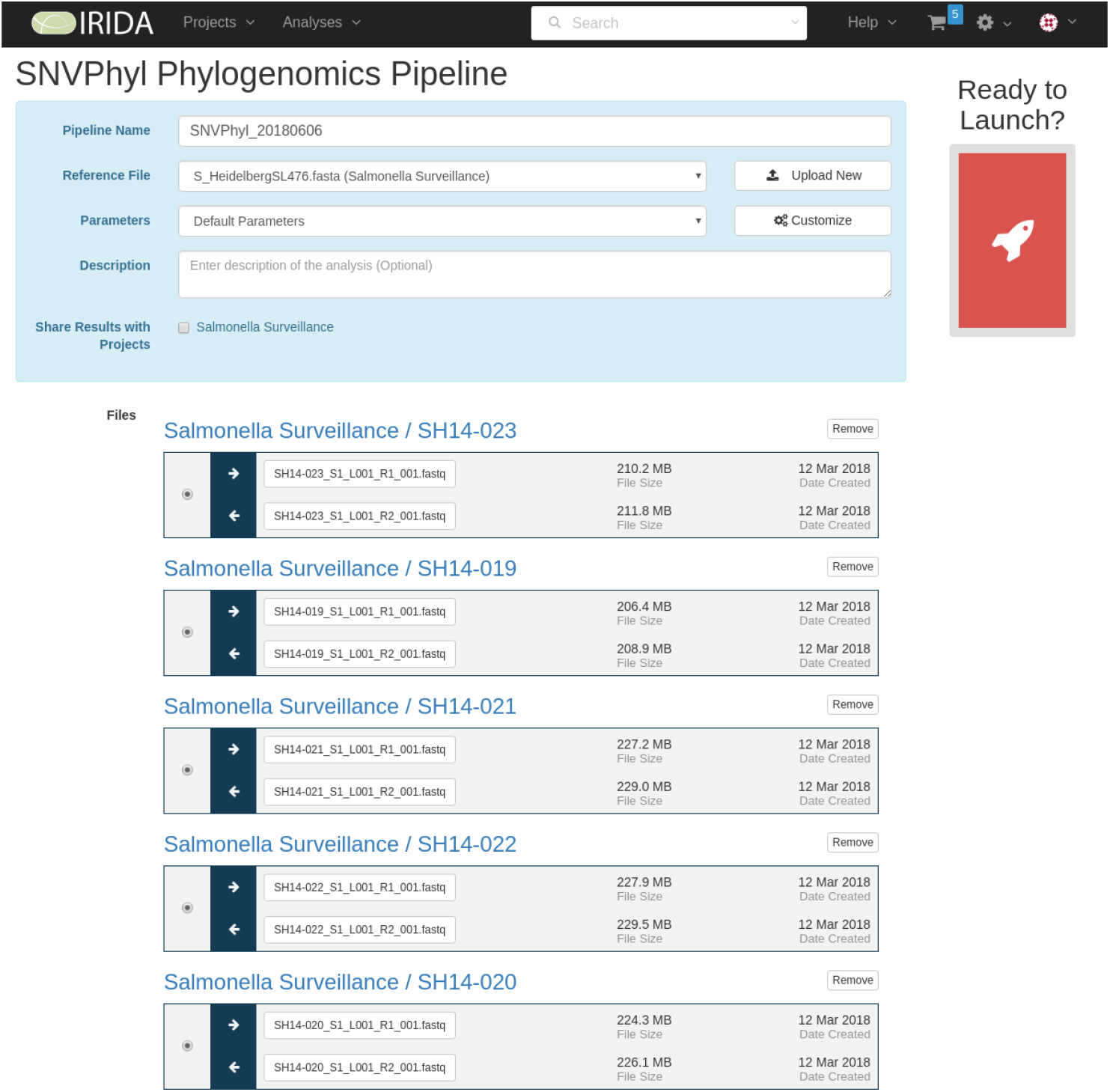
Pipeline launch. The pipeline launch page makes it as easy for users to launch advanced analysis tools. After adding samples to the cart and selecting a pipeline, users may adjust the parameters— optionally saving these adjustments for application to future pipelines. Clicking the “Launch” button initiates the pipeline.

One of IRIDA’s principal functions is to manage the processing of sequence data and metadata through its collection of analysis pipelines. IRIDA incorporates an internal instance of the Galaxy bioinformatics workflow manager (25) to assist in executing the analysis pipelines, which are implemented in IRIDA as Galaxy workflows (galaxyproject.org). Galaxy manages the distribution of analysis jobs on high-performance computing systems. IRIDA provides a computational superstructure on top of its internal Galaxy system to shield users from the complexity of running the analysis pipelines and to provide additional functionality not available in the Galaxy system such as automated processing of newly imported data, creation and selection of saved pipeline parameters, management of analysis results within a project, and incorporation of analysis outputs into the metadata tables contained within a project.

IRIDA provides several ways to export sequence data, metadata, and analysis results. Users can download their sequence data on to their local workstations through a web-browser. IRIDA also can export data to a locally installed “companion” installation of Galaxy. This feature is useful for running analytical pipelines that are not directly integrated within the IRIDA system. Advanced users with Linux workstations and having access to the IRIDA file system can export their sequence data to their user accounts using a command-line linking utility. Rather than copy the sequence data, both the Galaxy exporter and the command-line linking utility create symbolic links to the original sequence files avoiding unnecessary data duplication and ensuring that the integrity of the original data is preserved.

### User Management and Data Sharing

IRIDA is developed as a multi-user application, providing industry standard user authentication and user access controls. Authorization to access data and operations are controlled throughout the application including web interface and REST API. A number of roles are in place to control what a user can and cannot do. The roles fall into two categories: User and Project. Within the User category there are four roles. The first role - User - is intended for regular users; this role grants authority to create and manage projects. The Sequencer role provides the authority to upload data to any project. This role has no login privileges and cannot perform any operations nor access any data on the system outside of data upload. Manager roles have the same privileges as Users, and have additional authority to invite other Users to the system and to modify User details. The Admin role has full system privileges. Admins can access all projects and see all analyses. Roles in the Project category include Project Manager and Project Collaborator. Project Managers “own” the project data. They can import new sequence data and metadata, and can modify existing metadata. Project Managers also can update project settings, invite other users to the project, and share data to another project. Project Collaborators have authority to view sequencing data and metadata for specified projects, and can share data to other projects with limited privileges. They also can run analysis pipelines on data contained within a project. Project Collaborators are not authorized to modify any existing data within a project.

IRIDA uses a project-centric data sharing model and features the ability to share data with other users within the same installation or share data across different installations. Changing of permissions is permitted transiently or permanently by IRIDA Administrators, who can manage their User Groups in such a way as to alter single profiles or assign permissions in bulk to any number of users. Project Managers can share their project data with other users on the same system by simply adding them to the project. Sharing data across IRIDA instances is accomplished through “synchronized projects” and uses a host-client model. The host IRIDA instance provides connection privileges to the client instance. A user on the client instance connects to the host and associates a project on the host instance with a local project; the client instance then pulls the sequence data and metadata from the host project into a local project.

As the host project acquires new data, or modifies existing data, the client will automatically synchronize the data. Each IRIDA instance can synchronize any number of projects with any number of remote IRIDA instances, thus enabling users to create *ad hoc* data sharing networks. Project synchronization can be revoked at any time by the client or the host, although the data shared among IRIDA instances cannot be revoked once it has synchronized across instances.

IRIDA is also capable of submitting sequence data and metadata to NCBI’s sequence read archive (SRA), which is synchronized daily with EBI-EMBL’s ENA and Japan’s DDBJ. This functionality allows sequences to then be easily shared with global, centralized resources such as GenomeTrakr. Users select the datasets they wish to make public, the associated metadata they wish to publish, and the NCBI BioProject ID they have established prior to data deposition. IRIDA will then perform the data submission automatically. Once the data is received by NCBI, IRIDA retrieves and saves the accession numbers.

### Sequence Analysis

IRIDA provides the ability to analyze genomic sequence data and metadata using its collection of analysis pipelines. Newly imported sequence data can trigger the execution of certain analysis pipelines, such as genome assembly or *in silico* serotyping. Combined with the automated sequence Uploader Tool, this gives IRIDA the ability to automatically generate analysis results without any user intervention. Analysis results can be incorporated back into the project’s metadata and can be viewed in IRIDA as a line list. All results can be exported for use outside of IRIDA.

The sequence analysis pipelines and utilities currently included in IRIDA have been developed to serve the requirements of the Canadian public health system. We summarize them briefly in the following sections.

### Quality Control

IRIDA automatically runs FastQC (26) to perform a quality check on newly imported sequence reads. Results of FastQC (quality scores, read duplication, overrepresented sequences, *etc*.) are captured by IRIDA and presented as charts and tables. Additional quality checks, such as sequence coverage, are calculated for samples within projects. Users can set their own quality score cutoffs for each project. IRIDA flags files that fail quality control before they are included in an analysis, and flags analyses that fail to run for any reason. In-depth information for QC and analysis failures is provided as “drill down” pages.

### Genome Distance Calculation

IRIDA provides fast genome distance calculation through the RefSeqMasher pipeline (github.com/phac-nml/refseq_masher), which uses Mash (Mash v2.0+) (27) and a k-mer based Mash sketch database of 54,925 NCBI RefSeq genomes to enable users to quickly match the submitted isolate reads to the closest matching reference genomes. The RefSeq Masher pipeline can be used to select appropriate reference genomes for use with SNVPhyl or to identify possible contamination. In IRIDA, RefSeqMasher results are presented in tabular form, sorted by the closest matching genome.

### Single Nucleotide Variant Detection and Phylogenomic Inference

IRIDA uses SNVPhyl to detect single nucleotide variants (SNVs) contained in genomic sequence data, and applies these SNVs to infer a phylogenetic tree (28). Sequence reads generated from a collection of isolates are mapped against one user-selectable reference sequence using SMALT (http://www.sanger.ac.uk/resources/software/SMALT/), and high quality SNVs are detected from the resulting pileups using FreeBayes (29) and SAMtools/BCFtools (https://github.com/samtools/bcftools). The concatenated hqSNVs are organized into a multiple alignment, and a maximum likelihood tree is estimated from the alignment using PhyML (30). SNVPhyl provides a table of all identified SNVs and a SNV distance matrix. SNVPhyl can mask internally repeated sequences as well as regions of higher SNV density, indicating possible recombination. Additionally, users may mask user-selectable regions of the reference sequence, such as mobile elements.

### Assembly and Annotation

IRIDA provides an Assembly and Annotation pipeline, which enables users to assemble and annotate a single genome or a collection of genomes in batch. Paired-end reads are first merged using FLASh (31), and then passed to SPAdes (32) to perform a *de novo* assembly. Contigs returned by SPAdes are filtered to remove small and low-coverage contigs; quality-filtered contigs are then annotated by Prokka (33). Output files for each sample include assembly statistics, a list of contigs, and Prokka-generated annotations. Batch assembly and annotation enables the user to download a single output data package for all submitted samples.

### Serotype Prediction

IRIDA currently performs *Salmonella* serotype prediction with the *Salmonella In Silico* Typing Resource (SISTR), a validated bioinformatics platform for rapid *in silico* inference from draft *Salmonella* genome assemblies (34). SISTR performs highly-accurate serovar prediction based on genoserotyping through sequence analysis of the *Salmonella* O and H antigens, with additional refinement of predictions based on population structure context via cgMLST analysis and genomic distance calculation using MASH (9,34,35). Results generated by SISTR are then incorporated into the Sample metadata and can be conveniently viewed in a single table.

### Multi-Locus Sequence Typing

IRIDA facilitates cgMLST analysis through the integration of MentaLiST, a fast k-mer based MLST and cgMLST calculation engine enabling genotyping of bacterial samples directly from read data (36). MentaLiST’s ability to call alleles directly from raw sequence reads bypasses time-consuming assembly, and is specifically designed and implemented to handle large typing schemes (*i.e*. thousands of loci) while requiring minimal computational resources. A distance matrix can be constructed based on the Hamming distances between allelic profiles for each sample. The Bio.Phylo module from the Biopython library is used to calculate a neighbour-joining tree in Newick format (37). Within IRIDA, the Newick tree can be displayed using IRIDA’s advanced visualization system, and allelic profiles are provided in downloadable tabular form.

### Visualization and Reporting

IRIDA provides support for visualizing metadata and analysis results. Trees built by SNVPhyl or MentaLiST are viewable using a modified version of PhyloCanvas (phylocanvas.org) supporting real-time visualization of trees with thousands of taxa directly in the web browser (Fig. 5). Trees can be displayed in a variety of familiar layouts such as rectangular, circular, and radial views. Users can customize various tree attributes (e.g. colors, labels) and export trees to bitmap (PNG) or vector (SVG) formats.

**Figure 5.**
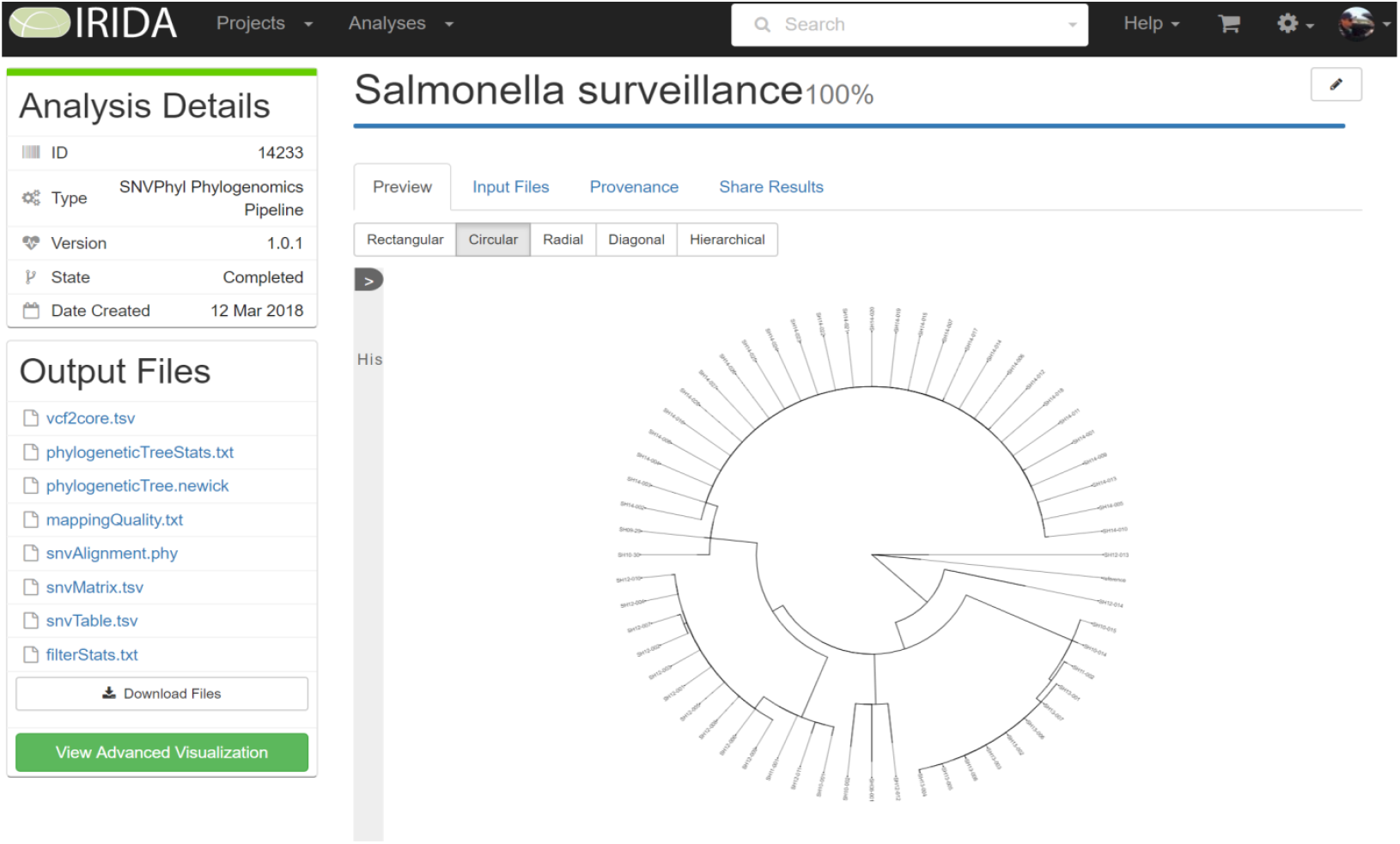
Example of a SNVPhyl phylogenomics analysis. Once an analysis from one of IRIDA’s different pipelines is completed, the results are stored in IRIDA and can be viewed within the browser, e.g. the SNVPhyl circular SNV-based tree presented here. A number of pipelines have custom visualization tools associated with them to quickly view results. All analysis results can be exported for further analysis in the user’s preferred external tools. Provenance information also can be viewed from this page, which displays all information about how the analysis pipeline was executed, including all tools, versions, and parameters that were used to build an analysis result.

Sample metadata contained within a Project can be viewed in an interactive tabular format, modeled on the “line list” format commonly used to coordinate outbreak investigations (Fig. 6). Columns corresponding to metadata categories can be rearranged or hidden to best present the relevant data for a given investigation. Users can save their customized table formats as templates, which can be applied to any number of projects. Metadata tables can be exported as a spreadsheet in Microsoft^®^ Excel or CSV format. IRIDA’s Advanced Phylogenomic Visualization tool integrates phylogenetic trees displayed by PhyloCanvas with epidemiological metadata. The integration of genomic relatedness information with epidemiological data can assist in outbreak investigations and enhances decision making by visualizing the covariation of genomic data and associated epidemiological, clinical, and biological trait data (Fig. 7).

**Figure 6.**
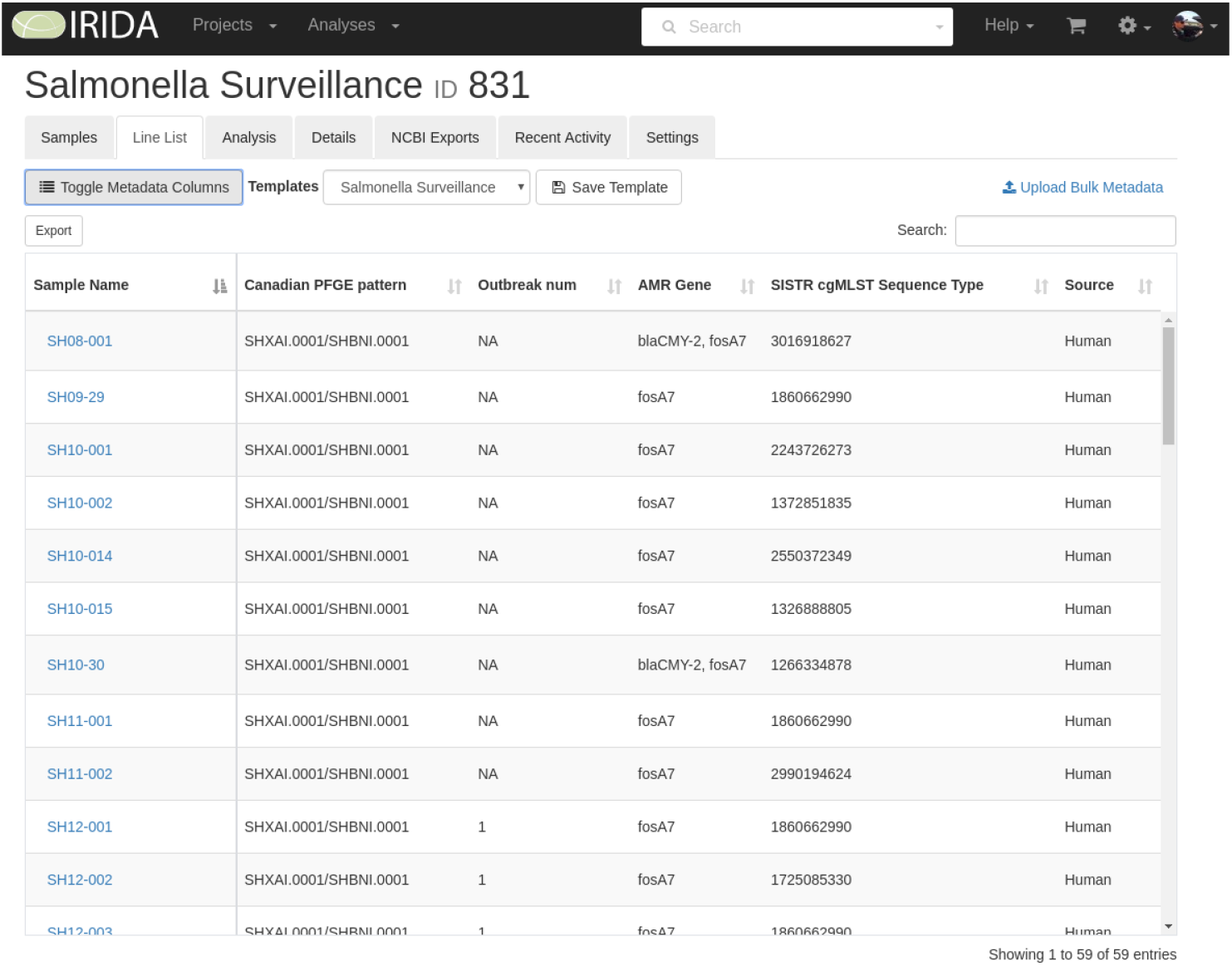
Example of line list displaying contextual information. IRIDA’s line list display provides users with a high-level view of all available contextual metadata for samples in a given project. This view allows users to sort, import, and export any terms available in their samples. Imports and exports of data from IRIDA’s metadata system are in Microsoft^®^ Excel or csv format, allowing users to easily update or share data outside of IRIDA. Visibility of columns in the table can be toggled on or off, and rearranged, enabling different projects or users to have customized views on the same datasets. These views can be saved as a metadata template, allowing users to quickly return to a given view. Metadata templates can be defined by project managers, configuring metadata terms for their project or investigation.

**Figure 7.**
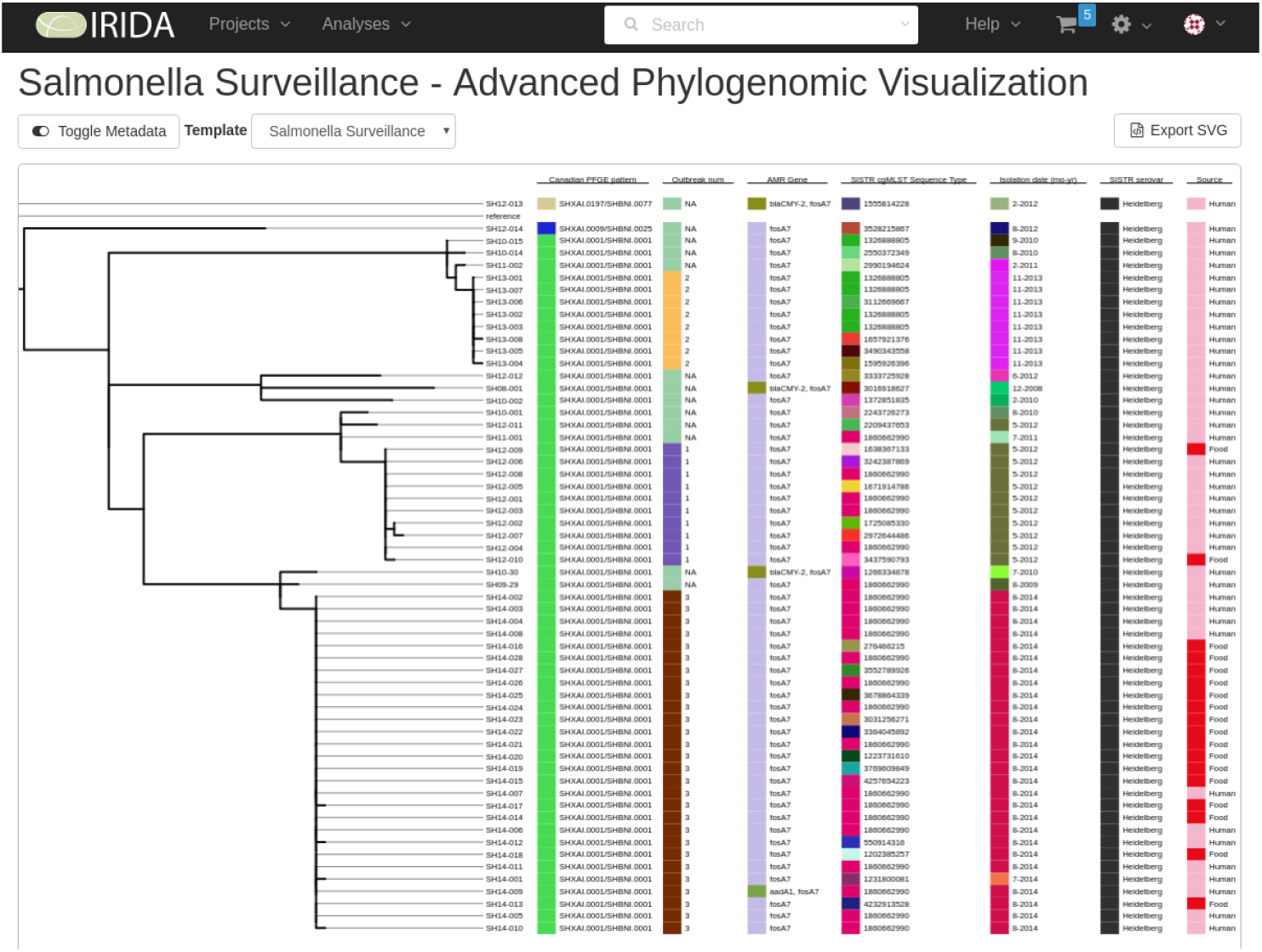
Example of integrated phylogenomic tree with contextual data. IRIDA’s Advanced Phylogenomic Visualization tool combines results of IRIDA’s SNVPhyl phylogenomic pipeline with the contextual metadata from the samples used to build the phylogenetic tree. This visualization tool aligns the metadata from IRIDA’s line list tool with the leaves of the tree. This example displays a phylogenetic tree of *Salmonella* isolates, labelled with attributes such as sample source, pulsed-field gel electrophoresis (PFGE) typing information, antimicrobial resistance genes, and cgMLST plus SISTR sequence type data. Metadata can be toggled similar to the line list tool, and metadata templates can be applied to the fields. Colour coding is displayed to assist users in grouping metadata terms. This visualization can be exported in SVG format.

IRIDA generates automated quality reports for all WGS data uploaded into the system using FastQC. Users may examine quality reports on a sample-by-sample basis, or they may examine reports in bulk across many samples - primarily used for sequencing coverage, with low-coverage samples flagged by IRIDA. Analysis pipelines produce differing reports - such as genome length and contig N50 for the assembly pipeline, or MLST results for MentaLiST - but all pipelines include a report on the input data files, tools, and parameters used to generate the results. Organism-specific reporting may be enabled on a project-by-project basis. Currently, this is only supported for *Salmonella* serotyping, generated using SISTR and conforming to the existing report format used within the Canadian Public Health Laboratory Network (CPHLN).

As most public health, clinical microbiology, and food safety laboratories have specific requirements for reporting the results of their analytical tests, IRIDA provides a REST API for external report-generating applications. One application to make use of the REST API is the IRIDA SISTR Results software (github.com/phac-nml/irida-sistr-results), which exports *Salmonella* serotyping information, in bulk to an Excel spreadsheet. IRIDA’s web interface also provides the ability to export metadata, genome sequence data, visualizations, and analysis results in standard file formats to facilitate integration into custom reporting systems that cannot directly interact with IRIDA via its REST API.

### Linkage to External Tools

IRIDA provides a REST-API which enables authorized external programs to interact with IRIDA and enhance its analytical capabilities. One such external program is GenGIS, a free and open-source bioinformatics application that allows users to combine digital map data with information about biological sequences collected from the environment from multiple sample sites (38). GenGIS provides a connector to IRIDA to download the results of phylogenetic analyses and geographic information stored within IRIDA, integrating this information into a phylogeographic map. Another external program making use of the IRIDA REST API is the BioNumerics software suite, facilitating PulseNet Canada’s surveillance activities.

An instance of IRIDA maintained by Simon Fraser University (Burnaby, Canada; sfu.irida.ca) also features integrated API access to IslandViewer (pathogenomics.sfu.ca/islandviewer/), a genomic island (GI) prediction and visualization tool (39). IslandViewer has been used for detection of GIs in *Listeria monocytogenes* isolates obtained from food, food processing environments and listeriosis patients in Canada and Switzerland, as well as the detection of *Streptococcus pneumoniae* isolates obtained from an outbreak in British Columbia (40,41). Results from IslandViewer analyses are returned to IRIDA in tab-delimited or GenBank formats, and include links to interactive circular and linear layouts of the results on the IslandViewer website.

## Methods

### Architecture

#### Development Technologies

IRIDA is developed as a Java servlet-based web application built using the Java Spring Framework. The web interface is built using AngularJS (angularjs.org), Bootstrap (getbootstrap.com), Webpack (webpack.js.org) and Thymeleaf (thymeleaf.org). IRIDA’s core database is versioned with Liquibase (liquibase.org) to ensure updates to the database are applied consistently during an upgrade. IRIDA uses the public GitHub software code repository for hosting, versioning, bug tracking, feature requests, task management, and software documentation (github.com/phac-nml/irida).

#### Analysis Pipelines

IRIDA uses the Galaxy workflow management system to carry out the execution of IRIDA’s analytical pipelines on high-performance computing systems (25). IRIDA prepares sequencing data, analysis parameters, and other workflow information, and provides these to Galaxy, which manages the execution of the pipeline – implemented as a Galaxy workflow. If installed on a computer cluster, Galaxy also will manage the distribution of analysis jobs across the cluster’s available resources. IRIDA communicates with Galaxy via its API (galaxyproject.org/develop/api/) and monitors the progress of the analysis as it proceeds through the pipeline. Upon completion, IRIDA retrieves the analysis results along with details about the pipeline such as component software versions and database versions.

IRIDA is designed to simplify the integration of additional pipelines by external developers. External developers can include pipelines not provided by IRIDA by building the pipeline as a Galaxy workflow, then integrating the pipeline into IRIDA via an internal API designed to expose the pipeline’s analysis parameters. IRIDA automatically generates web forms to present the analysis parameters to the user. The decoupling of the analysis pipelines from the IRIDA base system allows for easy modification using the Galaxy workflow editor. Thus, IRIDA’s analysis capabilities can be tailored to the needs of individual labs.

IRIDA provides a REST API to facilitate interoperability by external programs, including scripts, desktop applications, and web applications. The API supports authentication via OAuth2.0. Once authenticated, external applications can download sequence data, metadata, and analysis results for the projects in which they have been granted access. Existing Project, Sample, and File metadata can be modified and new metadata added, depending on the access privileges granted by IRIDA to the external application.

### Best Practices

IRIDA aims to comply with best practices for public health labs, clinical microbiology labs, and food regulatory authorities (21,24). Areas of particular focus include data security, data integrity, reproducibility, transparency, validation of software performance, and validation of analysis results.

#### Security

IRIDA has been designed with data security as a priority. IRIDA uses the Spring Security framework to control access to all data in IRIDA. Spring Security ensures that the roles assigned to users of the system have the correct authorization level to view or modify any data within IRIDA. Security controls are applied across all access points including the web interface and the REST API. The REST API uses the industry-standard OAuth2.0 protocol to delegate access to external applications. Access via the REST API can be granted or revoked on an application-by-application basis by IRIDA administrators. IRIDA enforces strong password policies including a configurable password expiry period.

#### Software Testing and Development

The IRIDA core development team follows a strict software development process to ensure the application performs as expected. All code written for IRIDA undergoes intensive review by the development team to ensure proper documentation and testing guidelines are followed. A comprehensive testing protocol, including unit tests and integration tests, is carried out to verify the integrity of the code base. User acceptance tests are conducted to verify the application is fit for use within the Canadian Public Health Laboratory Network. Before deployment, IRIDA must undergo testing in multiple environments, including a development environment, an integration testing environment, and a mock production environment.

#### Data Integrity

IRIDA implements the Java Hibernate Envers (hibernate.org/orm/envers) module for data auditability. All data created, modified, or deleted in IRIDA is audited with the timestamp, user, and tool which was used to modify the data. Reports can be generated for tracing the history of samples and sequencing information. Deleted or modified data can be recovered and restored to prior versions. All sequence data imported into IRIDA is stored in a read-only format in its underlying file system. IRIDA additionally supports reproducibility by capturing the provenance for all analyses performed by its integrated pipelines. Component software versions, parameters, database versions, and ancillary workflow information are recorded and provided as a viewable report within the web application, or downloadable as a JSON-formatted file. All versions of the pipelines are stored within IRIDA to allow users to re-run and reproduce their results if desired.

### Availability

IRIDA is open source software released under the Apache 2.0 license. Participation in developing IRIDA’s codebase by the research community is encouraged. IRIDA can be installed on workstations or high-performance computing clusters. IRIDA requires a Linux, Java, Tomcat, and MySQL/MariaDB environment for installation. Analyses require a configured instance of Galaxy. IRIDA is intended to be installed on a network-accessible filesystem for operation on a compute cluster. Alternatively, a pre-built VirtualBox virtual machine is also available for evaluation purposes.

#### Training and Support

All IRIDA software is supported by documentation and user guides available online (irida.corefacility.ca/documentation/). Technical support for installing and operating IRIDA and its associated pipelines comes in two forms. For agencies operating within Canada’s Public Health Laboratory Network, the Public Health Agency of Canada (PHAC) offers direct support through the IRIDA development team operating within the National Microbiology Laboratory in Winnipeg, Manitoba. For support outside the Canadian public health system, IRIDA requests are serviced through the open-source community support model (available at github.com/phac-nml/irida/issues).

## DISCUSSION

The bioinformatics skills and computational sophistication required to perform data analysis of large genomics data sets are currently beyond the reach of many public health, food safety, and clinical microbiology laboratories. Critical gaps still exist to integrate WGS into laboratory workflows. These gaps include the lack of laboratory capacity in decentralized healthcare systems to perform localized genomic data processing and analyses, and the lack of easy and automated methods to share data across jurisdictions without uploading to a centralized repository. The IRIDA platform reduces the expertise required for bioinformatics analysis, and provides technical solutions for multi-jurisdictional data sharing in accordance with jurisdictional data governance policies. Moreover, as an open-source platform, additional and customized functionalities can be added by third parties to address individual laboratory needs. The design principles of the IRIDA platform were developed to specifically address these critical gaps, as well as to better integrate bioinformatics into public health environments, to strengthen clinical and public health interfaces, and to facilitate local-to-global information exchange. A number of other bioinformatics tools and platforms have been developed to better enable laboratories to utilize genomics data, each of which may have different associated costs and best practices (Table 1). Although IRIDA is primarily designed to serve the needs of Canada’s multi-jurisdictional public health system, it can be used by anyone wishing to analyze microbial genomic sequence data, and instances have been installed in countries spanning four continents.

**Table 1.** Comparison of commonly used public health genomics software according to best practices. The IRIDA platform is compared with INNUENDO, WGSA, the Centre for Genomic Epidemiology (CGE), BioNumerics, SeqSphere, Nullarbor, COMPARE and NCBI public health genomics software, using cost and best practices criteria. Only IRIDA, INNUENDO and Nullarbor are considered completely open-source. Only IRIDA, SeqSphere and BioNumerics enable independent installations to perform private analyses in a decentralized system with data management and storage features, however work is underway to enable this functionality in INNUENDO. While many software packages and platforms record data provenance and parameters, some are predefined and cannot be changed by the user. All data created, modified, or deleted in IRIDA is audited with the timestamp, user, and tool that was used to modify the data. All software offer quality control checks. Only IRIDA software currently enables users to share data with other instances, as well as public repositories, in an automated fashion. Most non-commercial software is free; in contrast, the licenses for the commercial packages SeqSphere and BioNumerics carry significant costs.

| Criteria | IRIDA | INNUENDO | Center for Genomic Epidemiology | WGSA | Nullarbor | COMPARE | SeqSphere | BioNumerics | NCBI Pathogen Detection |
| --- | --- | --- | --- | --- | --- | --- | --- | --- | --- |
| Open-Source | ✓ | ✓ | Partially | ✗ | ✓ | Partially | Proprietary | Proprietary | Partially |
| Free | ✓ | ✓ | ✓ | ✓ | ✓ | ✓ | ✗ | ✗ | ✓ |
| Centralized/ Localized | Localized | Centralized | Centralized | Centralized | Localized | Centralized Data Hubs | Centralized (Star Topology) | Centralized (Star Topology) | Centralized |
| Interface | Web interface & REST-API | Web interface | Web interface | Web interface | Command line/ Web interface | Web interface | Graphical User Interface | Graphical User Interface | Web interface |
| Private Data Storage & Management | Private Projects and Samples | Private Projects and Samples; accessible by all users of each installation | Temporary storage; Operates as service only | Private Projects and Assemblies | Private Projects and Samples | Consortia Data Hubs (contain Private Projects and Samples) | Private Projects and Samples | Private Projects and Samples | Public repository with embargo periods for submitted sequences |
| Auditability | Provenance & customizable parameters captured and archived; metadata changes recorded and tracked | Predefined / Custom installation workflows could enable provenance & customizable parameter capture | Predefined parameters | Predefined parameters | Provenance & customizable parameters captured and archived | Project-dependent parameters | Provenance & customizable parameters captured and archived | Provenance & customizable parameters captured and archived | Limited documentation |
| Quality Control Checks | ✓ | ✓ | ✓ | ✓ | ✓ | ✓ | ✓ | ✓ | ✓ |
| Data Sharing | Between IRIDA instances and with NCBI SRA | Between users of the same installation / Manual | ✗ | Assemblies can be shared with public WGSA collection | Manual | Between members of a Data Hub | Central Repository | Central Repository | Public repository |
| Publication/ Website | irida.ca | sites.google.co | (57) | wgsa.net | github.com/tsee | (20) | ridom.com/seqsph | applied- | ncbi.nlm.nih.gov/p |
|  |  | m/site/theinnu<br>endoproject | genomicepidemi<br>ology.org |  | mann/nullarbor | compare-europe.eu | ere/ | maths.com/bionum<br>erics | athogens |

### IRIDA is an analysis environment for large-scale microbial genomics

IRIDA’s web interface guides users through microbial genomics data analysis, from creating projects, uploading WGS samples, and analyzing the samples, to visualizing the analysis results and sharing these results with other users. IRIDA uses a “shopping cart” model to assist users in selecting data for analysis pipelines. IRIDA presents the progress of each pipeline and completed results in the web interface; results can also be exported for use in third-party applications. IRIDA records detailed information for each analysis pipeline and generates a provenance report for auditing purposes.

#### IRIDA performs analyses of sequence data and metadata with validated pipelines and integrated visualizations

Currently, IRIDA provides validated, user-friendly, fit-for-purpose, pipelines for analyses and integration of WGS and metadata. The SNVPhyl pipeline has been validated for analysis of foodborne disease outbreaks, and correctly distinguishes outbreak-related isolates from non-outbreak isolates across a range of parameters, sequencing data qualities, and in the presence of contaminating sequence data (28,42–44). Since 2010, SNVPhyl has been used for analysis by hundreds of public health analysts at Canada’s National Microbiology Laboratory to support research and provincial laboratory services. The pipeline is currently being used for outbreak investigations and has been validated as part of a suite of tools used by PulseNet Canada for all genomic epidemiological investigations of foodborne outbreaks since 2012.

In addition to SNV-based methods, IRIDA provides functionality for performing MLST. Current MLST approaches that extend classical MLST schemes to include an organism’s core genome (cgMLST) or whole genome (wgMLST) have been validated for surveillance of foodborne disease, and are quickly being adopted by surveillance programs such as PulseNet International, and used for a variety of nosocomial, zoonotic, and tuberculosis outbreaks (45–49). IRIDA has adopted the fast, accurate, and computationally efficient MentaLiST system as its main pipeline for cgMLST-based analysis. The MentaLiST pipeline generates MLST allelic calls directly from read data, avoiding slow and computationally expensive genome assembly. MentaLiST’s allele calling concordance with other popular genome-scale MLST programs is greater than 99% (36).

The computational resources required to characterize and compare genomes can be immense, with billions of bases generated per sequencing run and petabytes of publicly available sequence data (50,51). Modern data reduction techniques are extremely valuable. Mash is one such general-purpose toolkit that applies the MinHash technique to efficiently reduce large sequence read sets to compressed sketch representations (27). IRIDA offers a Mash-based pipeline called RefSeq Masher to quickly identify the closest genome matches, select appropriate reference genomes for SNVPhyl, as well as identify possible contamination.

*In silico* serotyping of *Salmonella* is an alternative approach to classical serotyping that identifies *Salmonella* serovars directly from genomic sequence data, which has the benefit of being cheaper, higher throughput, and automatable. IRIDA includes in its pipeline collection the *Salmonella In Silico* Typing Resource (SISTR), a bioinformatics application for rapidly performing simultaneous *in silico* analyses for several leading subtyping methods on draft *Salmonella* genome assemblies. SISTR has been extensively validated, yielding accuracies of ~95%, the highest among serotype prediction tools (9). SISTR is currently being used to generate serotype predictions for all genomes submitted to EnteroBase, the largest global repository of *Salmonella* WGS data (enterobase.warwick.ac.uk/). SISTR has generated predictions for the 30% of *Salmonella* genomes that have been deposited at NCBI with missing serovar information (9). Implementation of SISTR also has led to the phasing out of antigen-based serotyping at Canada’s National Microbiology Laboratory, which, as of May 2018, has moved to employ WGS as the primary means for characterizing *Salmonella* isolates from national surveillance programs (9).

IRIDA generates dynamic line lists based on uploaded or manually entered sample metadata and other contextual information. The line list is the primary tool used by epidemiologists to collect and organize preliminary information on cases under investigation. Epidemiologists can analyze and integrate contextual information with genomics data in the same platform—critical for the rapid identification of outbreaks, when the alternative is the difficult and time-consuming process of moving large WGS files between analytical platforms. This dynamic feature enhances decision making and can be used to identify isolate clusters fitting the criteria for triggering outbreak investigations, establishing case definitions, and developing response protocols.

#### IRIDA enables localized analyses, as well as data sharing

IRIDA instances behave as independent data management and analysis environments, with fine-grained data-access security controls. Through IRIDA’s decentralized framework, data sharing across IRIDA instances can be fine-tuned through User and Project permissions; these can be customized according to data governance guidelines, bypassing the need to transmit sensitive information by traditional means such as fax, phone, mail or email. Furthermore, sequence data, metadata, and analysis results can be exported for offline use by authorized users. IRIDA also provides native support for depositing sequence data and metadata to the public archives, supporting global data sharing and complementing centralized resources.

#### IRIDA is contributing to the creation and adoption of community data standards

The FAIR (Findability, Accessibility, Interoperability, and Reusability) Guiding Principles prescribe best practices for “metadata” (or contextual data) knowledge integration and reuse by the community (52). To make contextual information capture for infectious disease and food safety investigations more FAIR, the IRIDA team has created two new open-source and publicly available ontologies, the Genomic Epidemiology Ontology (GenEpiO) and the Food Ontology (FoodOn), in collaboration with two new international consortia (www.genepio.org; www.foodon.org). Ontologies are hierarchies of well-defined and standardized vocabulary interconnected by logical relationships (53). These logical interconnections provide a layer of intelligence to query engines, making ontologies much more powerful than simple flat lists of terms (54). The IRIDA team continues to work toward the integration of ontology-derived metadata specifications and ontology terms within the platform. Uptake of interoperable ontologies between different public health genomics platforms and tools will better enable the communication of results and information, which will be a crucial step towards realizing a truly responsive global infectious disease surveillance system.

### IRIDA Deployment and Uptake

IRIDA is the official bioinformatics platform for public health genomics within the Public Health Agency of Canada. It has been in use since 2016 by Canada’s provincial and national public health laboratories for genomic investigations of foodborne disease outbreaks, as part of PulseNet Canada’s foodborne disease surveillance activities. Instances of IRIDA have been installed across the globe (e.g. the United States, Switzerland, Singapore, South Africa, Italy), including a demonstration version currently available in the Genomics Virtual Lab toolkit (55). Potential users wishing to test the functionality of the IRIDA platform can do so in a number of different ways: as a public demo instance maintained for demonstration (hosted by Simon Fraser University; sfu.irida.ca), private instance installation (requires a working and configured install of Galaxy), or install of an IRIDA virtual machine (downloadable, fully-configured virtual appliance).

### Future Development

Future IRIDA developments include ontology integration, enhancements of privacy controls and data exchange mechanisms, the creation of tools for modeling gene transmission dynamics as well as expert systems for genome assembly and annotation. Future pipeline development and integration will aim to decouple individual pipelines from the platform, facilitating a plugin-style architecture to allow developers to individually package their pipelines. Ongoing efforts include the integration of IslandViewer as a core module to enable viewing of genome, annotation and genomic islands in a visually dynamic way, from the whole genome down to individual genes and enabling side-by-side comparison with other publicly available genomes (39). The IRIDA consortium of developers will also work towards the creation of tools to enable the export of data to other external web applications through provided APIs, for example, the Comprehensive Antimicrobial Resistance Database (CARD) and its Resistance Gene Identifier (RGI) providing antimicrobial resistance and virulence determinant prediction (56). We are also working to enhance IRIDA’s deployability, maintainability, portability, and support for cloud computing.

## CONCLUSION

The IRIDA platform (irida.ca) is a user-friendly, distributed, open-source bioinformatics and analytical web platform developed to support real-time infectious disease outbreak investigations using whole genome sequencing data. IRIDA strives to shield public health, regulatory and clinical microbiology laboratory personnel from the computational challenges of genomics big data by providing solutions for data storage, management, analyses, and data sharing, to better incorporate WGS technology into routine operations. The functionality and control over data provided by IRIDA can help to overcome the many pervasive challenges for real-time global infectious disease surveillance, investigation and control, resulting in faster responses, and ultimately, better clinical outcomes and improved public health.

## ACKNOWLEDGEMENTS

The authors gratefully acknowledge the IRIDA Consortium (Marsha Taylor, Eleni Galanis, Linda Hoang, Natalie Prystajecky, Kim Macdonald, Lynn Schriml, David Aanensen, Michel Dumontier and William Klimke), as well as Celine Nadon, Aleisha Reimer, Lorelee Tschetter, Natalie Knox, Shaun Tyler, Cameron Sieffert, Sukhdeep Sidhu, Shane Thiessen, Paul Williams the GRDI-FWS Consortium, and SFU Research Computing, for their critical feedback and support.

## ETHICAL STATEMENT

There is no human or animal work in this study.

## Abbreviations

AMR: antimicrobial resistance
API: application programming interface
CARD: Comprehensive Antibiotic Resistance Database
cgMLST: core genome multi-locus sequence typing
CPHLN: Canadian Public Health Laboratory Network
DDBJ: DNA Database of Japan
EMBL-EBI: European Molecular Biology Laboratory - European Bioinformatics Institute
Gi: genomic islands
GMI: Global Microbial Identifier
INSDC: International Nucleotide Sequence Data Consortium
IRIDA: Integrated Rapid Infectious Disease Analysis
JSON: JavaScript Object Notation
MLST: multi-locus sequence typing
NCBI: National Center for Biotechnology Information
PFGE: pulsed-field gel electrophoresis
PHAC: Public Health Agency of Canada
PNG: Portable Network Graphics
REST: Representational State Transfer
QC: quality control
SNV: single nucleotide variant
SRA: Sequence Read Archive
SVG: Scalable Vector Graphics
wgMLST: whole genome multi-locus sequence typing
WGS: whole genome sequencing.

